# Prion safety laboratory swipe test

**DOI:** 10.1101/2024.11.15.623841

**Authors:** Sara M. Simmons, Qi Yuan, Jason C. Bartz

## Abstract

Transmission of prion disease has occurred from contaminated neurosurgical tools, transplant materials and from occupational exposure to prion contaminated laboratory tools. Prions cause disease by the templated misfolding of the normal cellular form of the prion protein, PrP^C^, into the misfolded and pathogenic form PrP^Sc^ and are invariably fatal. Reducing iatrogenic and occupational prion transmission is challenging. First, prions can bind to and persist on surfaces for long periods of time. Second, prions are highly resistant to inactivation. Given this, surfaces can retain infectivity for long periods of time following ineffective decontamination. Not only can this pose a potential occupational risk for prion laboratory workers but could potentially cross contaminate laboratory experiments utilizing sensitive prion amplification techniques. The protocol described here for a prion safety laboratory swipe test includes steps for the identification and documentation of high traffic laboratory areas, recommended swabbing controls to ensure validity of results, steps to identify proper responses to positive surface swabbing sites, representative results from prion swipe testing, as well as potential artifactual results. Overall, the prion safety laboratory swipe test can be implemented as part of a broader prion safety program, to assess decontamination of surfaces, monitor common spaces for prion contamination, and implement the documentation of prion decontamination status.

**SUMMARY:** A method to assess commonly used areas in laboratory settings for prion contamination and effective decontamination is lacking. The protocol described here provides key fundamentals for implementing a laboratory prion safety swipe test that can easily be modified to meet the individual needs of specific laboratories.

## INTRODUCTION

Prion diseases are invariably fatal neurodegenerative diseases with no known treatment or cure. Prion diseases are caused by PrP^Sc^, the misfolded and pathogenic form of the normal cellular form of the prion protein, PrP^C^ [1–5]. Prion diseases are known to affect humans and several other animal species. One human prion disease, Creutzfeldt-Jakob Disease, CJD, has three known etiologies: sporadic, inherited and acquired. Acquired CJD can occur as a result of accidental transmission (iatrogenic and occupational) and is thought to be the cause of Kuru in the Fore people of Papua New Guinea [6].

Prion transmission has been associated with prion contaminated medical devices and transplant materials [7–17]. Iatrogenic transmission of CJD can occur via blood, tissue or from prion contaminated surfaces [18–20]. For example, iatrogenic CJD can develop in patients following an electroencephalogram with electrodes previously used on an individual in the preclinical stage of CJD who then later succumbed to CJD [21]. More recent laboratory based occupational transmission has also occurred where a laboratory worker contracted prion disease via a skin puncture with forceps used to handle brain slices from an animal infected with sheep adapted BSE [22, 23]. Such transmission scenarios could occur within clinical, laboratory and diagnostic laboratory settings where prion samples are handled.

Prions resist common disinfection techniques and can persist and remain infectious on surfaces for extended periods of time [24–29]. Common disinfection techniques such as the use of ethanol, phenolic cleaners, hydrogen peroxide, various forms of radiation, and formaldehyde are inadequate for the inactivation of prions, allowing surfaces to retain infectivity [30–37]. These characteristics contribute to the transmission of prions during iatrogenic and occupational exposure.

Methods for detection of environmental prions have only recently been developed. An environmental swabbing method coupled with RT-QuIC can assess residual prion infectivity from environmental surfaces as well as common laboratory surfaces following ineffective disinfection [38–40]. Here we describe how this technique can be incorporated into a broader prion safety program. Overall, this method can allow for the monitoring of laboratory dependent disinfection protocols, the investigation and proper documentation of contamination status which can help ensure the validity of experiments by minimizing cross contamination, the assessment of shared use spaces for prion contamination and allows for directional retraining of personnel based on commonly contaminated areas.

## PROTOCOL

**Figure 1.**
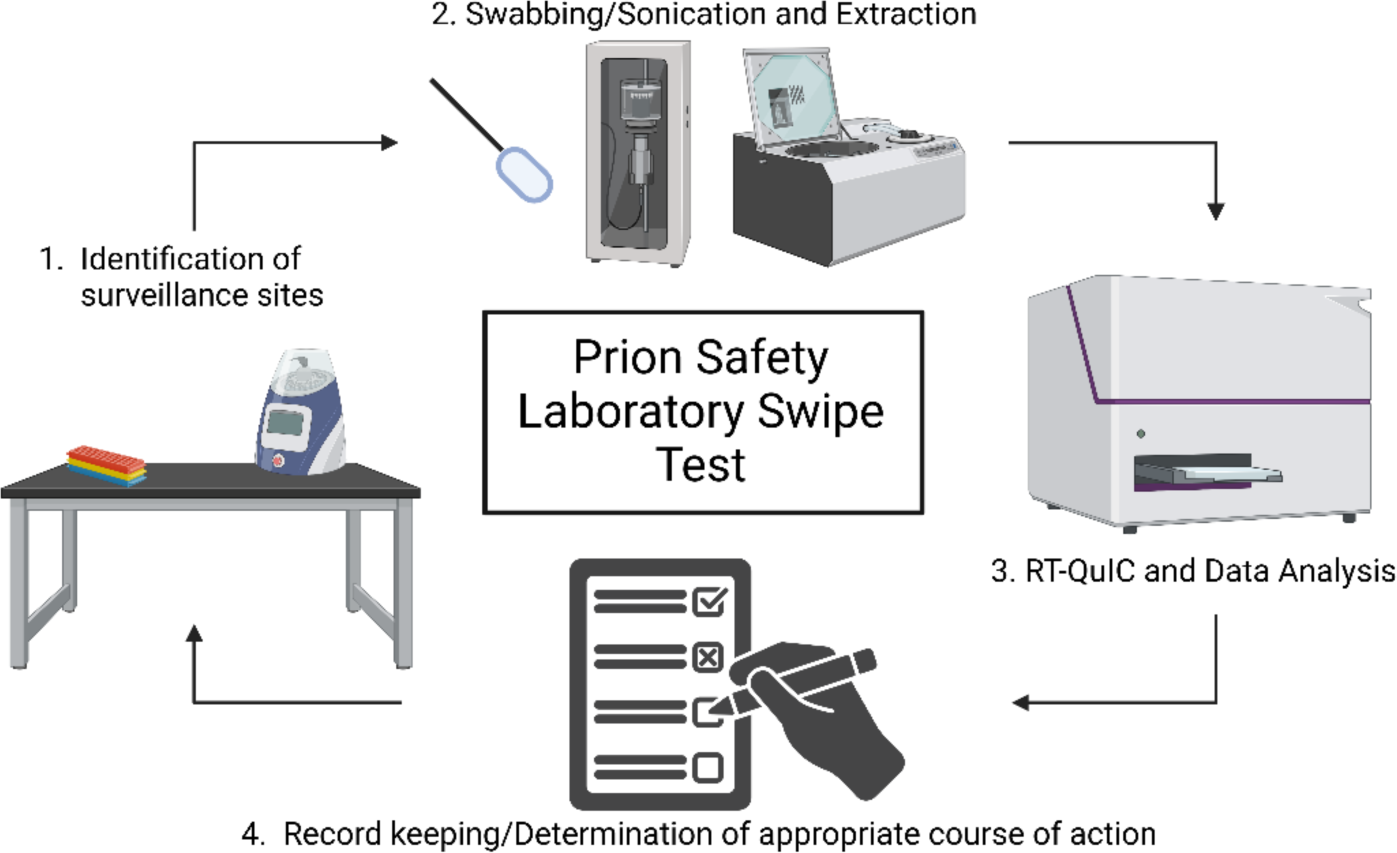
Schematic overview of prion safety laboratory swipe test.

### I. Selection of swabbing sites and preparation for surface swabbing

1.1: Identify and label appropriate surveillance areas for swabbing. A template is provided for documentation and reference (Figure 6 and S1)
1.2: Retrieve foam tipped swabs from storage in an area free of prion contamination.
1.3: Prepare a squeeze bottle with MQ H_2_O for use in step 2.1.

### II. Positive and negative control swab preparation

2.1: Prepare dilutions for positive and negative swab controls using relevant control types. Example: For positive control: prepare a 1% dilution of prion infected brain homogenate in DPBS. For negative control: prepare a 0.1% dilution of uninfected brain homogenate from the same species as the positive control in DPBS.
2.2: Using clean gloves, retrieve the appropriate number of swabs from the clean packaging and place the handle side down into a tube rack, taking care to space out foam swabs so that the tips are not contacting other swabs or surfaces. For each control sample and swabbing location three swabs should be prepared. Gloves should be changed between each sample.
2.3: Holding a clean foam swab by the handle, apply 50 μl of respective positive and negative control samples to foam swab tips. Make sure to apply to both sides of the swab tip to ensure complete absorption. Note: always change gloves prior to removing a clean swab from the tube rack.
2.4: Using scissors, cut off the excess handle of the swab (approximately ½ of the length) and place the swab into the preloaded microcentrifuge tube with the foam tip portion pointed down. The foam tip should be submerged in the DPBS and the handle should be cut adequately to allow for the lid of the microcentrifuge tube to close completely.
2.5: Continue applying control samples, after changing gloves, until all samples have been applied.

### III. Surface swabbing

3.1: Holding a clean foam tip swab by the handle, prewet the foam tip with MQ H_2_O and shake off the excess. Place the moistened foam tip of the swab onto the area chosen for surveillance and swab the area back and forth, approximately ten times, while simultaneously rotating the tip of the swab on the surface.
3.2: Using scissors, cut off the excess handle of the swab (approximately ½ of the length) and place the swab into the preloaded microcentrifuge tube with the foam tip portion pointed down. The foam tip should be submerged in the DPBS and the handle should be cut adequately to allow for the lid of the microcentrifuge tube to close completely.
3.3: Discard gloves and place new gloves on before each respective swabbing site to minimize the probability of cross contamination.
3.4: Method should be repeated until all areas chosen for surveillance have been swabbed.

### IV. Swab extraction and vacuum concentration

4.1: Place microcentrifuge tubes into circular tube rack. Place microfuge tube rack into the cup horn sonicator water bath. Ensure the foam swabs in DPBS within the microcentrifuge tubes are below the surface of the water in the cup horn (the handle portion does not have to be submerged). Apply the following settings: 15 seconds total run time (5 seconds on, 5 seconds off) at approximately 75-85 watts.
4.2: Following sonication, centrifuge the tubes for approximately 15 seconds to collect DPBS in the bottom of the tube prior to transfer.
4.3: Using a P1000 pipette set to 250 µl, carefully collect all liquid (swab extract) from the bottom of the microcentrifuge tube and transfer into the corresponding microcentrifuge tube from the second, empty prelabeled set. Use the pipette tip to squeeze any excess liquid from the foam tip. Discard empty tubes containing swabs.
4.4: Turn on the vacuum concentrator 30 minutes prior to use. Set the following settings: Temperature 45⁰C: Heat time: 15 minutes, Run time: 2 hours, Vacuum: 5.1.
4.5: Place the microcentrifuge tubes, containing swab extracts, into the vacuum concentrator, ensuring the tubes are placed in a balanced manner and all tube caps are open.
4.6: Upon cycle completion, ensure that samples are completely concentrated (only the pellet remains). Pellets are stored at −80°C until utilized for RT-QuIC. Note: In some instances, additional vacuum concentration time may be required to ensure complete concentration. Remove all concentrated samples, leaving only the tubes that still contain liquid. Rebalance the remaining tubes within the concentrator and run at additional 1-hour increments until samples are completely concentrated.

### V. Preparation of swabbing controls for use in RT-QuIC

Note: The RT-QuIC controls should be performed prior to the assay of environmental swab extracts to ensure that contamination has not been introduced during the swabbing, extraction or concentration procedures.
Note: For example layouts, see figures 3 and 4.
5.1: Prepare an appropriate negative RT-QuIC plate control (e.g., uninfected brain homogenate) by diluting to appropriate concentration in tissue dilution solution (N2-0.1%SDS/PBS). Prepare a positive plate control dilution by diluting prion-infected brain homogenate to a dilution of 10^−3^ in tissue dilution solution.
5.2.: Remove previously stored control swab extract pellets from −80°C and resuspend each with 50 µl of MQ H_2_O by pipetting up and down approximately 10 times, followed by vortexing briefly. Allow samples to sit at room temperature while performing the remaining steps.
5.3: Load 2 µl of specified plate controls which should include negative plate controls (tissue dilution solution alone and uninfected brain homogenate) and a positive plate control (infected brain homogenate).
5.4: Load 2 µl of each positive and negative swab extract technical replicate into a minimum of four replicate RT-QuIC wells.
Note: If negative swabbing controls exhibit RT-QuIC seeding above a laboratory’s standard for a positive determination, this would indicate that contamination had been introduced during the swabbing, extraction and concentration process and that the swabbing experiment should be performed again

### VI. Preparation of samples for use in RT-QuIC

6.1: Prepare an appropriate negative plate control (ex: uninfected brain homogenate, uninfected lymph node, etc.) by diluting to appropriate concentration in tissue dilution solution (N2-0.1%SDS/PBS). Prepare positive plate controls by diluting infected brain homogenate to a dilution of 10^−3^ in tissue dilution solution.
6.2.: Remove previously stored swab extract pellets from −80°C and resuspend with 20-50 µl of tissue dilution solution (depending on suspected amount of contamination) by pipetting up and down approximately ten times, followed by vortexing briefly. Allow samples to sit while performing the remaining steps.
6.3: Load 2 µl of specified plate controls which should include negative plate controls (tissue dilution solution alone and uninfected brain homogenate) and a positive plate control (infected brain homogenate).
6.4: Load 2 µl of each swab extract into four replicate wells.

### VII. RT-QuIC analysis and results

7.1: Perform RT-QuIC according to individual laboratory protocols (Optional protocols: Yuan et al., 2022; Orru et al., 2024).
7.2: Parameters for determining positivity of a sample should be determined by each lab (see representative results, Figure 5).
7.3: Record results in the provided table and take appropriate action based on laboratory best practices (Figure 7 and S2).

## REPRESENTATIVE RESULTS

I. Written description of positive and negative results (including positive/negative plate and swab controls).

Negative control swabs are included in the surveillance swabbing to monitor for potential prion contamination that could be introduced during the swabbing, extraction and concentration process. The first RT-QuIC plate performed for a given monthly surveillance should include the positive and negative swab controls. Successful negative controls fail to cross the positive fluorescence threshold (Figure 2, Panel A). This result would indicate that contamination had not been introduced during the experimental procedures. Successful positive control swab extracts would exhibit positive seeding in all replicate wells for a given sample. The inclusion of a positive control dilution series allows for the determination of the sensitivity of prion detection for a given experiment (Figure 2, Panel A).

**Figure 2.**
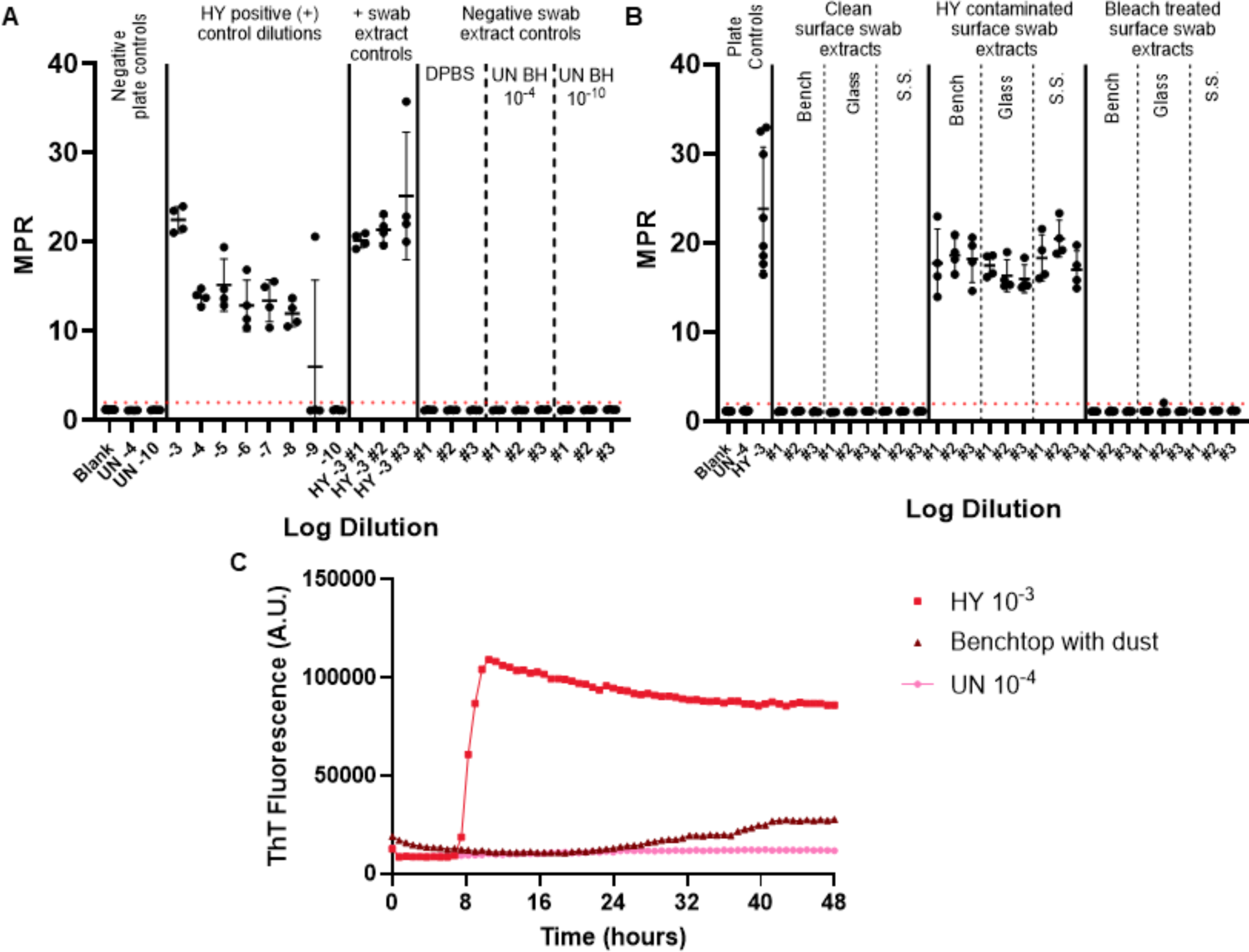
Representative surface swabbing experiment. A: Surface swab control plate containing negative swab extract controls in triplicate (DPBS, uninfected hamster brain homogenate 10^−4^ and 10^−10^) and positive swab extract controls in triplicate (HY brain homogenate 10^−3^). B: Representative swab extracts for benchtop, glass and stainless-steel surfaces that were swabbed prior to contamination, following contamination with HY 10^−3^, and following bleach treatment of contaminated surfaces. C: Fluorescence tracing comparison of uninfected brain homogenate 10^−4^, HY brain homogenate 10^−3^ and the swab extract from benchtop coated with a fine film of dust. Negative plate controls for panels A and B, include a blank (tissue dilution solution) and uninfected hamster brain homogenate 10^−4^. Panel B includes the addition of uninfected brain homogenate 10^−4^ and the positive plate control of HY brain homogenate 10^−3^. A positive fluorescence threshold (illustrated by red dashed line) was determined to be at 2. The maxpoint ratio (MPR) reported is the maximum fluorescence divided by the initial fluorescence reading obtained by the plate reader. Each point represents one technical well replicate for a given sample type. The mean and standard deviation are presented.

**Figure 3.**
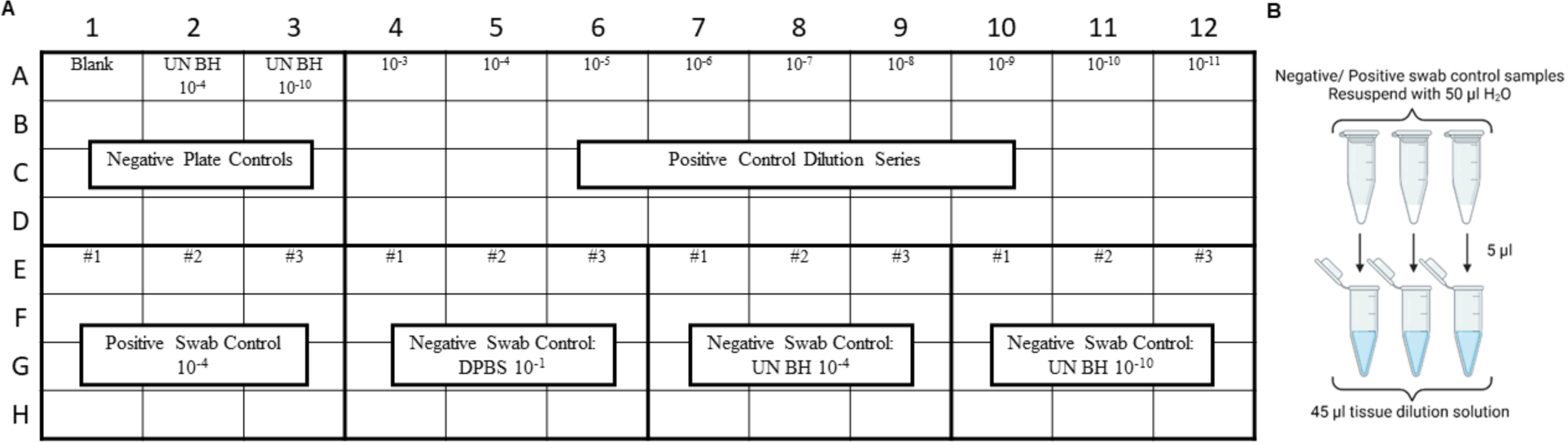
Sample experiment design for swab controls included prion safety laboratory swipe test. A. Sample layout for negative and positive swab extract controls B. Negative and positive swab extract controls should be resuspended with 50 μl of H_2_O. A 10-fold dilution of the resuspended swab extract should be generated by diluting into tissue dilution solution.

**Figure 4.**
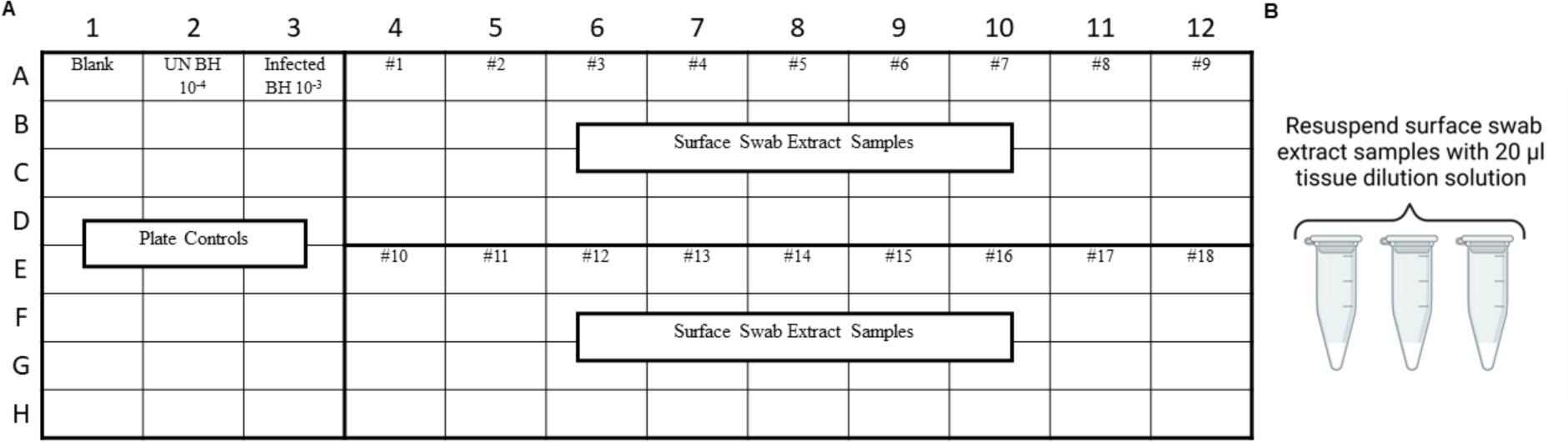
Sample experiment design for surface swab extracts from prion safety laboratory swipe test. A. Sample layout for surface swab extracts B. Surface swab extract samples should be resuspended with 20 μl of tissue dilution solution.

**Figure 5.**
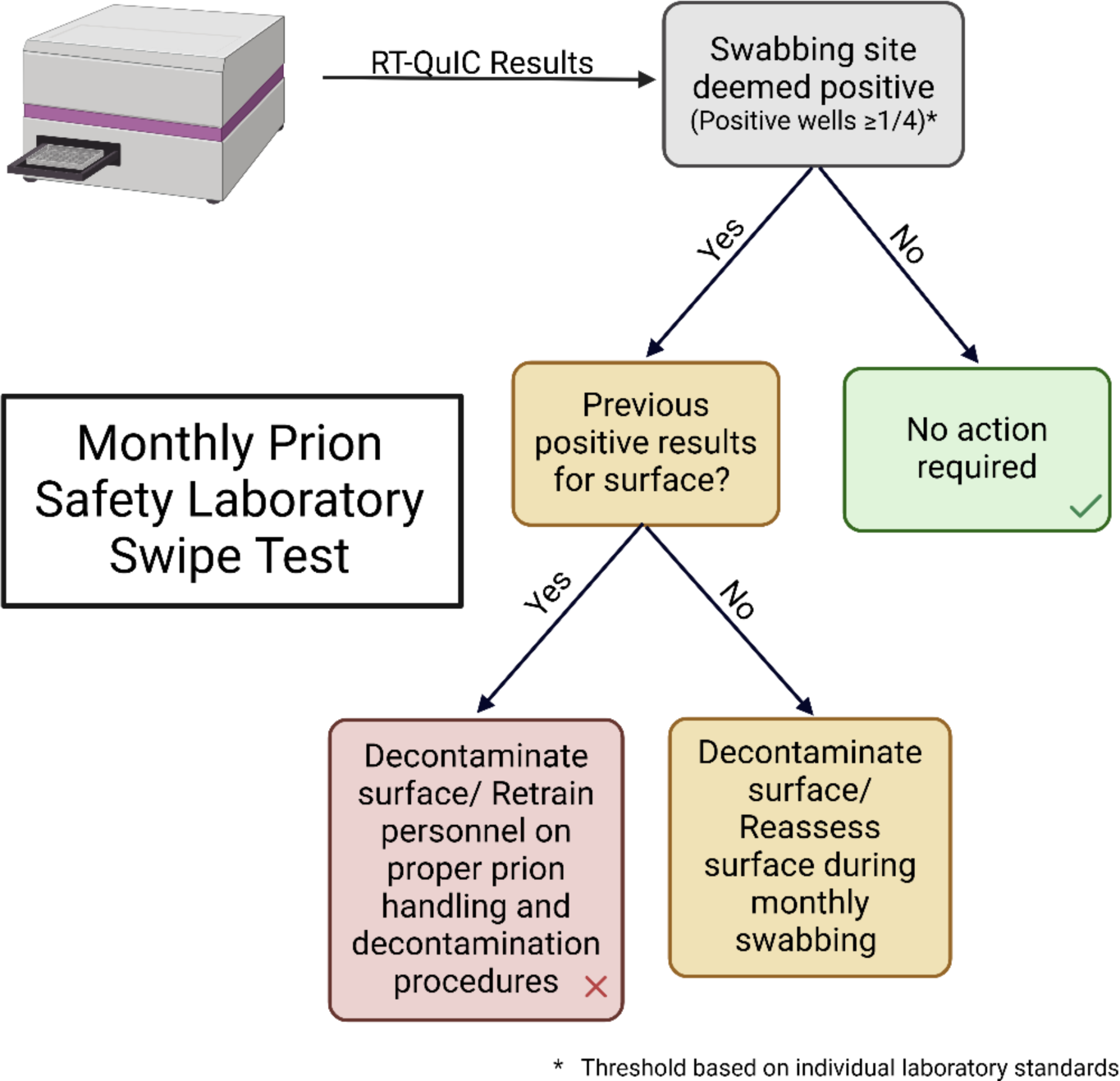
Monthly prion safety laboratory swipe test: flowchart for interpreting results and determining appropriate response.

**Figure 6.**
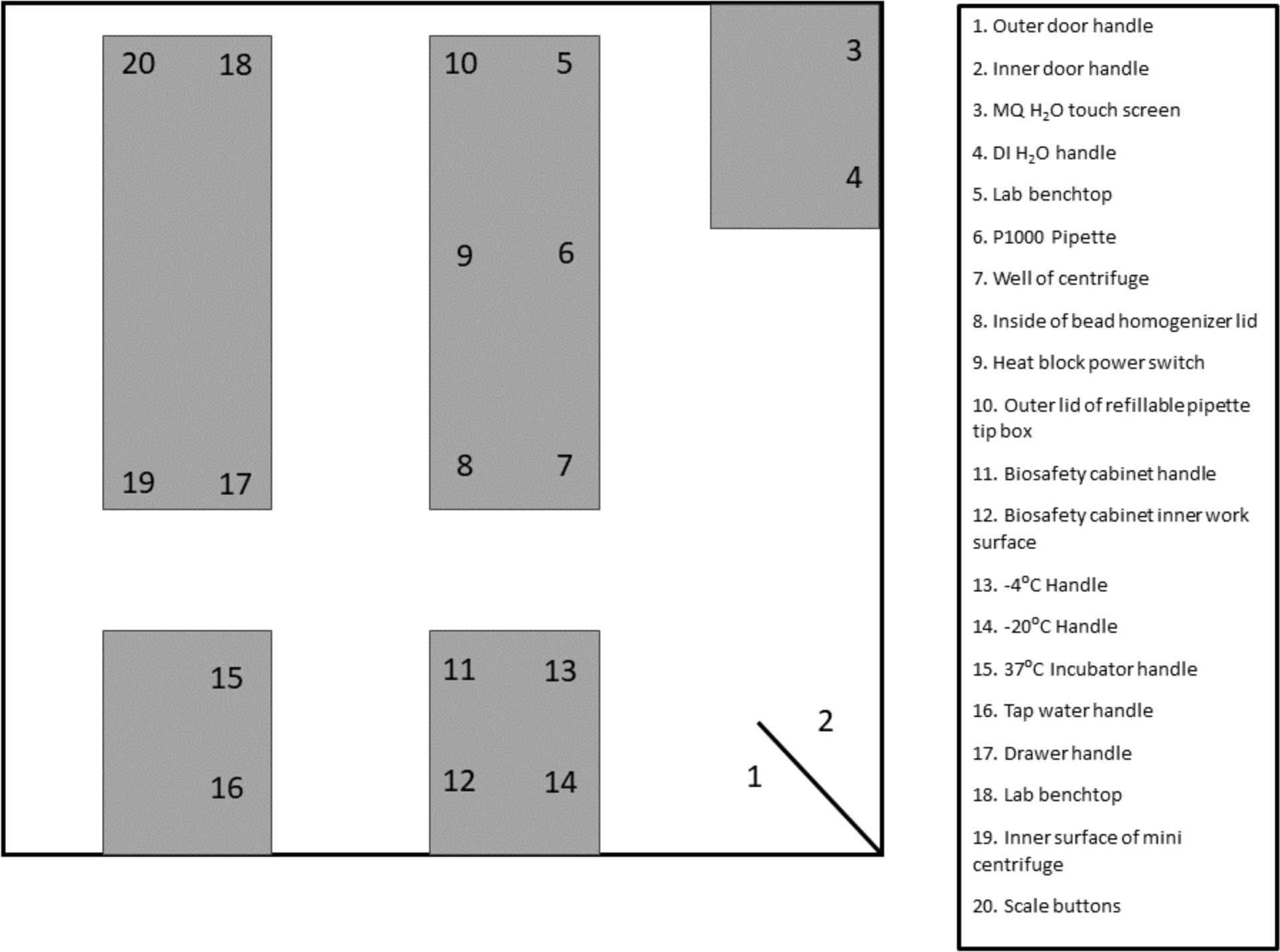
Sample swabbing site layout.

**Figure 7.**
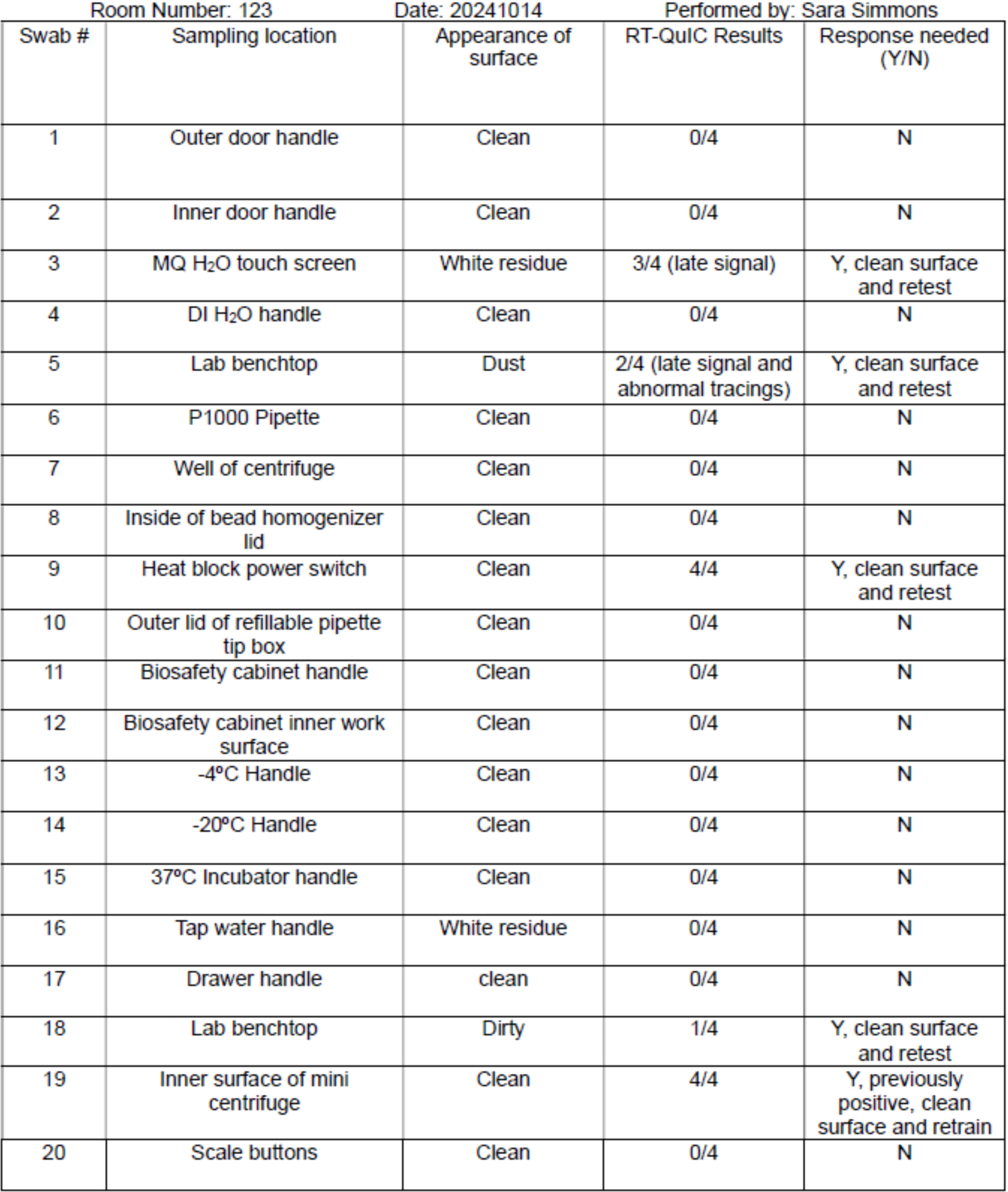
Sample swabbing site monthly documentation form.

When examining surface swab extract samples, surfaces that fail to show seeding above the predetermined positive fluorescence threshold can be considered to be prion negative (Figure 2, Panel B). Conversely, prion contaminated surface swab extracts will show seeding capabilities above the positive fluorescence threshold, although the MPR and time to fluorescence can vary compared to the included positive plate control (Figure 2, Panel B).The ability of the method to assess adequate disinfection is highlighted by the bleach treated prion contaminated surfaces, which now fail to seed RT-QuIC (Figure 2, Panel B).

Importantly, while our laboratory defines a positive sample as a sample that passes the set positive fluorescence threshold in at least half of the replicate wells, it is necessary for each laboratory to set their own standards. The rate of amyloid formation (RAF) and time to fluorescence can also be used to help establish laboratory specific thresholds for positivity.

Our laboratory has observed surface artifact results that pass the positive fluorescence threshold, but with altered kinetic curves and after an abnormally long time to fluorescence when compared to positive control samples (Figure 2, Panel C). These results should be cautiously interpreted as they may be caused by the presence of dust or residual chemicals present on a surface. These findings highlight the necessity of general laboratory cleanliness, as well as criteria for differentiating between true positive and false positive seeding.

## Supplemental Files

**Supplemental figure 1:** Swabbing site layout template.

**Supplemental figure 2:** Swabbing site monthly documentation form.

## DISCUSSION

The described prion safety swabbing method can be used to enhance existing prion safety measures. This method can monitor prion laboratory spaces and equipment, as well as shared laboratory spaces for potential prion contamination. Importantly, this method can be adapted to test laboratory specific disinfection techniques to verify decontamination of prion contaminated surfaces. As various prion strains display different sensitivities to disinfection techniques, this method can confirm that these techniques are effective for current laboratory experiments.

One limitation for this protocol is the potential generation of erroneous RT-QuIC results. Given the sensitive nature of RT-QuIC, the presence of residual detergents, salt and other substances on laboratory surfaces can impact the outcome of the reaction. Overall, given this observation, it is advantageous that surfaces remain free of substances that can interfere with RT-QuIC, such as: residual salt, detergents and dust.

A key benefit of this method for laboratories researching prions, is the ability to minimize contamination that may impact the results of sensitive amplification assays. Necropsy tools are commonly disinfected, prior to being reused for future necropsies. The described swabbing method provides a method to assess necropsy tools for residual prion infectivity that may affect downstream results. This can provide an additional level of rigor to experiments to exclude the possibility that prion detection in tissues was not due to contamination from the necropsy tools.

## Supporting information

Materials list

## ACKNOWLEDGMENTS

The work was supported by a grant from the Creutzfeldt Jacob Disease Foundation. The funders had no role in study design, data collection and interpretation, or the decision to submit the work for publication.

