## Supplemental Figure 1 for "Prion safety laboratory swipe test"

### Prion Safety Surveillance Layout:

**Instructions:** Draw layout of desired swabbing survey area and number the corresponding relevant equipment and work areas for swabbing.

**Suggested Areas:** High traffic work areas (benchtop, drawer handles, biosafety cabinets and pipettes), high traffic equipment (centrifuge, bead homogenizer, commonly used buttons/latches), door handles (doors to laboratory space, doors to freezers).

**Surface Number and Description:**
