## Supplemental Figure 2 for "Prion safety laboratory swipe test"

Room Number: ______ Date: ____________________________ Performed by: ____________________________

| Swab # | Sampling location | Appearance of surface | RT-QuIC Results | Response needed (Y/N) |
|---|---|---|---|---|
| 1 | | | | |
| 2 | | | | |
| 3 | | | | |
| 4 | | | | |
| 5 | | | | |
| 6 | | | | |
| 7 | | | | |
| 8 | | | | |
| 9 | | | | |
| 10 | | | | |
| 11 | | | | |
| 12 | | | | |
| 13 | | | | |
| 14 | | | | |
| 15 | | | | |
| 16 | | | | |
| 17 | | | | |
| 18 | | | | |
| 19 | | | | |
| 20 | | | | |
